# Modeling SIV kinetics supports that cytotoxic response drives natural control and unravels heterogeneous populations of infected cells

**DOI:** 10.1101/2020.01.19.911594

**Authors:** V. Madelain, C. Passaes, A. Millet, V. Avettand-Fenoel, R. Djidjou-Demasse, N. Dereuddre-Bosquet, R. Le Grand, C. Rouzioux, B. Vaslin, A. Saez-Cirion, J. Guedj

**Affiliations:** IAME, UMR 1137, INSERM, Université Paris Diderot, Sorbonne Paris Cité Paris, France; Institut Pasteur, Unité HIV, Inflammation et Persistance, Paris, France; CEA, Université Paris Sud, INSERM U1184, Immunology of Viral Infections and Autoimmune Diseases (IMVA), IDMIT Department / IBFJ, 92265 Fontenay-aux-Roses, France; Université Paris-Descartes, Sorbonne Paris-Cité, Faculté de Médecine, EA 7327, Paris, France; UMR 1065, INRA, Villenave d’Ornon F-33882, France

## Abstract

SIVmac_251_-infected Mauritius cynomolgus macaques presenting a M6 MHC haplotype or challenged with a low inoculum dose by mucosal route are models for natural HIV control. Here we characterized by modeling the dynamics of plasma SIV-RNA and of SIV-DNA in blood cells of 16 macaques of the ANRS SIC study.

SIV-RNA kinetics was best fitted using a model where the cytotoxic immune response progressively mounted up and reduced actively infected cells half-life (t_1/2_) from 5.5 days early on to about 0.3 days. The model predicted that the control was achieved in animals able to mount an effective immune response within three months, and this was corroborated by the longitudinal analysis of the CD8^+^ T-cell antiviral activity measured *ex vivo*. The control of SIV-RNA was accompanied in parallel by a slow and biphasic decline of SIV-DNA. This unravels the presence of at least two compartments of non-actively infected cells that are not rapidly eliminated by the immune system, one with a rapid turnover rate (t_1/2_=5.1 days) and predominant as long as SIV-RNA levels are still large, and one with a slow turnover (t_1/2_=118 days) consistent with the half-life of memory T-cells, and only visible when control is achieved,.

In summary, our analysis suggests that the establishment of an efficient CD8^+^ T-cell response in the first three months of the infection, and that progressively increases overtime is key to achieve SIV-RNA control in this model. Frequent SIV-DNA quantifications allowed identifying that most cells infected after viral peak have a short t_1/2_ but do not contribute significantly to viral production.

**One sentence summary:** Modeling viral dynamics in SIV natural controller macaques predicts that viral control is primarily driven by the capability to establish an efficient cytotoxic response and the viral decline during control unravels distinct compartments of infected cells.

## Introduction

Although, the vast majority of patients infected with HIV-1 progress to AIDS in about 10-15 years in absence of antiretroviral therapy *(1)*, a small proportion of patients are able to naturally maintain low levels of HIV-1 RNA plasma viral load *(2)*. These so called HIV controllers (HIC) represent less than 1% of HIV infected individuals *(1, 2)*, but they evidence that a control of HIV infection without antiretroviral therapy can be achieved. It has been reported that efficient CD8^+^ T-cell responses are strongly associated with HIV/SIV natural control *(3, 4)*. Furthermore, genetic factors, in particular major histocompatibility complex (MHC) alleles HLA-B*57 and HLA-B*27 *(1)*, are enriched in the cohorts of HIV controllers. Yet, the determinants of the establishment of viral control are difficult to characterize and up to now failed to be triggered in progressor patients *(5)*. Definitely, research is limited by the low prevalence of natural controllers, a number of confounding factors and the retrospective design of most studies, which limits the access to key information about viral dynamics during early stages of HIV infection in these patients *(6)*.

In this context, relevant non-human primate (NHP) models of SIV infection offer an opportunity to identify more in depth the mechanism of viral control *(7, 8)*. Asian macaques, particularly rhesus and cynomolgus infected with simian immunodeficiency virus (SIV) strains such as SIVmac239 and SIVmac251 are among the most popular models of SIV pathogenesis, as they closely reproduce the natural history of HIV infection as observed in patients, with an acute primary infection followed by a chronic phase, eventually evolving to AIDS in progressor animals *(9)*. Similar to humans, some MHC haplotypes/alleles, notably Mafa M6 in cynomolgus *(10)* or Mamu A*01, B*08 and B*17 in rhesus *(11–13)*, are associated with natural control of SIV infection. Moreover, infection through mucosal route with low inoculum dose was also found to improve the chance of viral control in cynomolgus macaques *(8)*, allowing to assess the mechanisms of control independent of a favorable genetic background.

Another advantage of NHP models is to allow a close monitoring of relevant markers over time that can be used in complement to plasma SIV-RNA in modelling studies. The quantification of total HIV/SIV-DNA in blood *(14)* has been shown to be predictive of disease progression and response to antiretroviral therapy, although its accuracy in estimating the size of viral reservoir remains debatable *(15)*. Indeed, total HIV/SIV-DNA reflects the global frequency of infected cells in peripheral blood but encompasses different forms of virus persistence, in particular linear and episomal unintegrated genomes, as well as replication competent and defective integrated genomes *(14)*. These forms may not contribute to virion production *(16)*, but may nonetheless have an impact on virus pathogenesis. In addition to the diversity of viral forms, the HIV reservoir encompasses heterogeneous infected cell populations (naïve, memory, and effector CD4^+^ T-cells, macrophages) *(17)*, with different lifespans and ability to expand through cell division. It was recently proposed that the majority of the infected circulating cells are clonally expanded, with provirus of archival origin *(18, 19)*, however the contribution of each cell population to viral reservoir and its association with an efficient immune response in the setting of natural control remain unclear.

Modeling viral kinetics after HIV/SIV infections has been the subject of numerous studies and has been instrumental to better understand viral pathogenesis and the efficacy of antiretroviral drug regimens *(20, 21)*. Over time, these mathematical models became more and more complex with the aim of characterizing not only the dynamics of viral replication, but also to better understand the development and evolution of efficient immune responses. Understanding the immune mechanisms associated with natural and post-treatment HIV control is of great relevance to the field. A recent study described a model that predicts immune responses and latent reservoir sizes needed for a patient to maintain viremia control after treatment interruption *(22)*. Despite the insights on HIV control gained through modelling, the prediction of immune responses and viral reservoir decay in the context of natural HIV/SIV control are yet to be developed.

Therefore, the aim of this study was to develop a model attempting to predict the determinants of a natural HIV/SIV control. In addition, by comparing the early kinetics of SIV-DNA in controllers and viremic animals, we aimed at bringing new insights on the characteristics of different cell compartments that contribute to HIV/SIV reservoirs.

## Results

### Viral metrics

In order to develop a model to predict the mechanisms associated with natural control of HIV/SIV infections and to characterize the different cell compartments contributing to HIV/SIV reservoirs, we took advantage of the ANRS SIC study *(23)*. Briefly, we analyzed the kinetics of SIV-RNA and SIV-DNA of cynomolgus macaques naturally controlling (n=12) and non-controlling (n=4) SIV infection, monitored for virologic and immunologic parameters along 18 months.

Kinetics of SIV-RNA and SIV-DNA loads are described in *(23)* (see also Figure S1). In brief, all macaques showed a similar pattern during acute infection, with a median SIV-RNA at peak of 6.2 (min-max: [5.13; 7.09]) log cp/mL and a median time to peak of 14 days ([11-17]). In contrast, median SIV-DNA at peak was equal to 3.7 log cp/mL ([2.7; 4.6]) and occurred slightly later at day 21 post-infection ([15; 36]). From peak to day 168 post-infection, SIV-RNA declined with a median slope of −0.038 ([−0.061; −0.015]) log cp/mL/day, compared to −0.011 ([−0.023; 0.0032]) log cp/mL/day for SIV-DNA. Consequently, (log) SIV-RNA/SIV-DNA ratio declined over time with a slope of −0.0133 /day ([−0.022; 0.00049]), reflecting that SIV-RNA decay was much faster than SIV-DNA decay (Table S1, Figure S1).

Finally, predictions could be confronted to experimental data obtained to characterize high quality CD8^+^ T-cell responses, as measured by the capacity of *ex vivo* SIV-specific CD8^+^ T-cells to eliminate autologous infected CD4^+^ T-cells (CD8 antiviral activity). Overall, the measurements of CD8 antiviral activity demonstrated a large inter-individual variability, with peak values ranging from 0.44 to 3.95 log ng/mL p27 antigen decrease, and a time to peak ranging from 15 to 567 days post-infection. The slope of SIV-RNA decay was significantly faster in those who subsequently achieved control below 400 copies of RNA per mL than in those with higher viremia, then determined as viremic macaques (p=0.013), but there was no difference in early SIV-DNA or CD8 antiviral activity kinetics between controller and viremic animals (Table S1).

### Immune response model

Based on previous results obtained for the characterization of HIV/SIV infections, (3, 5, 9, 11), we considered an adaptation of the target cell limited model accounting for the cellular cytotoxic immune response. In order to find the best parameterization of the model, we identified 9 candidate models from the literature (Table S2). Detailed information is given in the Material and Methods section.

The selected model among the nine candidate models was adapted from *(22)* and is given by the following set of ODEs:

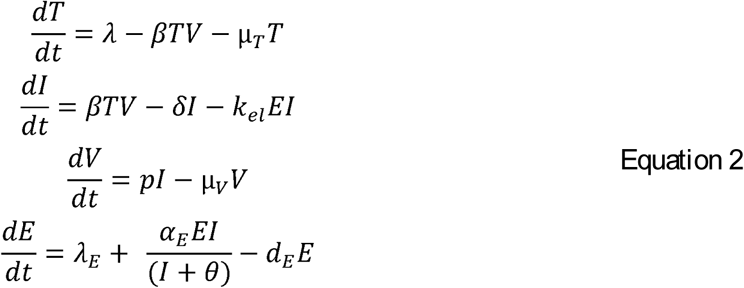

Whereas T, I and V are target cells, actively producing infected cells and free virions, respectively, E represents the effectors of the cytotoxic immune response. In this model, cells are infected and become actively producing with a rate β, then they are naturally lost with rate δ, while they can also be lost with a rate k_el_*E representing the effect of the cytotoxic effector response in targeting and eliminating actively producing cells. Importantly, in this model the cytotoxic effector immune response growth depends on the number of actively producing cells but saturate, and θ represents the number of infected cells needed to induce half of the maximal proliferation rate, and thus represents how fast the immune system “senses” and mounts its response. Eventually, we included a cytotoxic effector elimination rate, d_E_, which represents all potential cumulative mechanisms that could lead to a progressive loss of efficiency of the immune response, such as apoptosis or immune exhaustion.

### Final model

Next, we included SIV-DNA assuming one infected cell contains one copy of viral DNA, but the model could not fit jointly SIV-RNA and SIV-DNA (Figure S2). This is due to the fact that this model assumes that all infected cells are actively producing and hence predict that SIV-DNA, as it is the case for SIV-RNA, should rapidly decline after peak, which is in contradiction with the data observed (Table S1).

In order to capture the discrepancy between SIV-RNA and SIV-DNA, we extended the model assuming that SIV-DNA containing cells could also account for short-lived cells that are infected but cannot produce virus and/or long-lived cells that are infected and may potentially become actively producing after a latency period (Figure 1). We found that a model incorporating the three infected cell compartments, actively producing infected cells (I), short-lived non-actively producing actively infected cells (S) and long-lived non-actively producing infected cells (L), provided the best fit to the data (Table S3 and Figure 2). Of note, the model predicts that the observed SIV-DNA decline with a biphasic kinetic after the peak. We verified that this prediction was not artefactual by fitting SIV-DNA kinetics after the peak with a simple mono or bi-exponential function, and we found that a bi-exponential function was in fact statistically supported in 6 out of 16 macaques (Figure S3).

**Figure 1:**
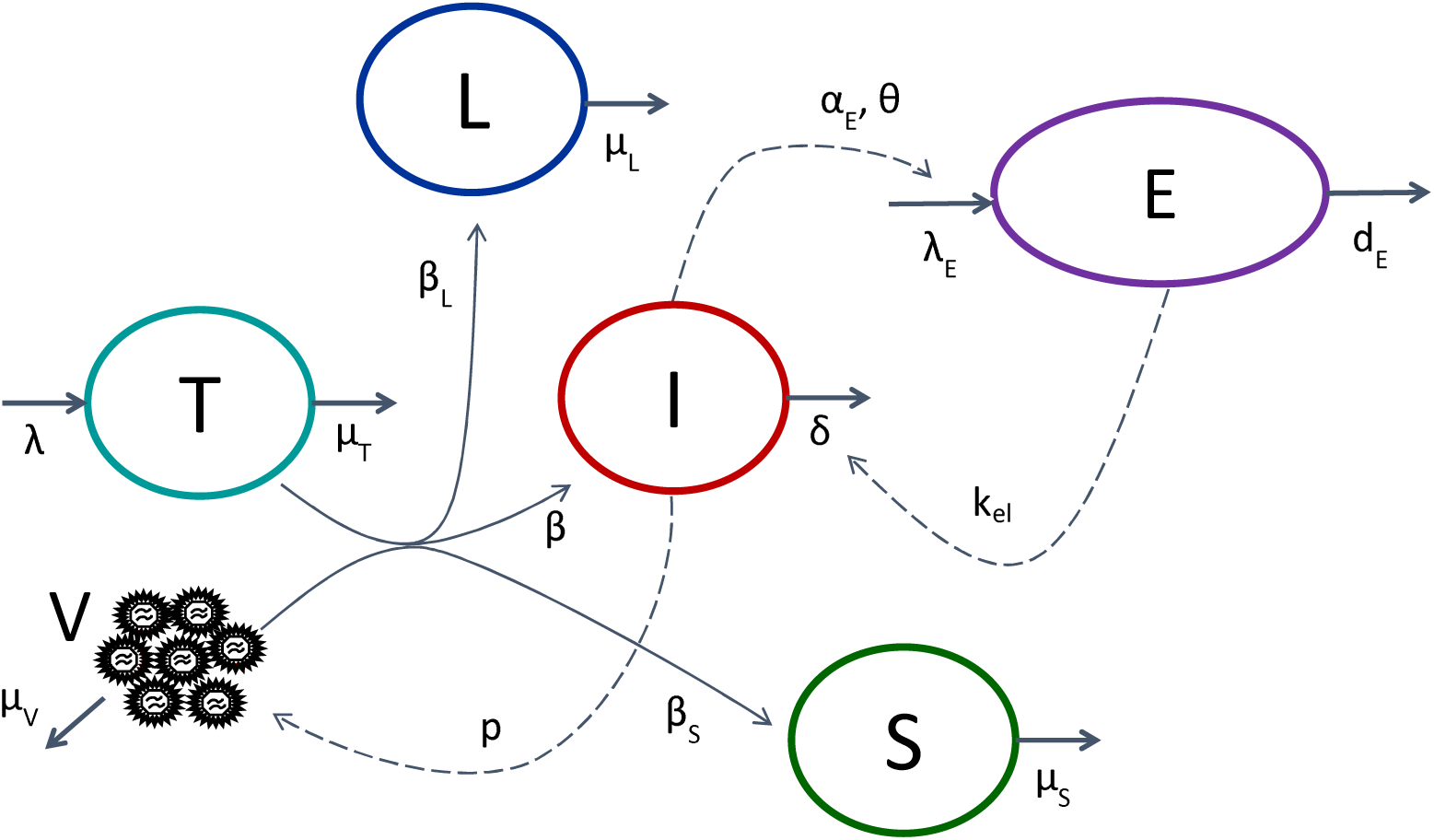
Viral kinetic model used to fit SIV-RNA and SIV-DNA kinetics. T, V and I account for target cells, virions and actively infected cells (eg, producing virus), respectively. S and L account for non-actively infected cells (i.e. not producting virus) that are either short-lived or long-lived, respectively, while E represent the immune cytotoxic response, which is stimulated by actively infected cells.

**Figure 2:**
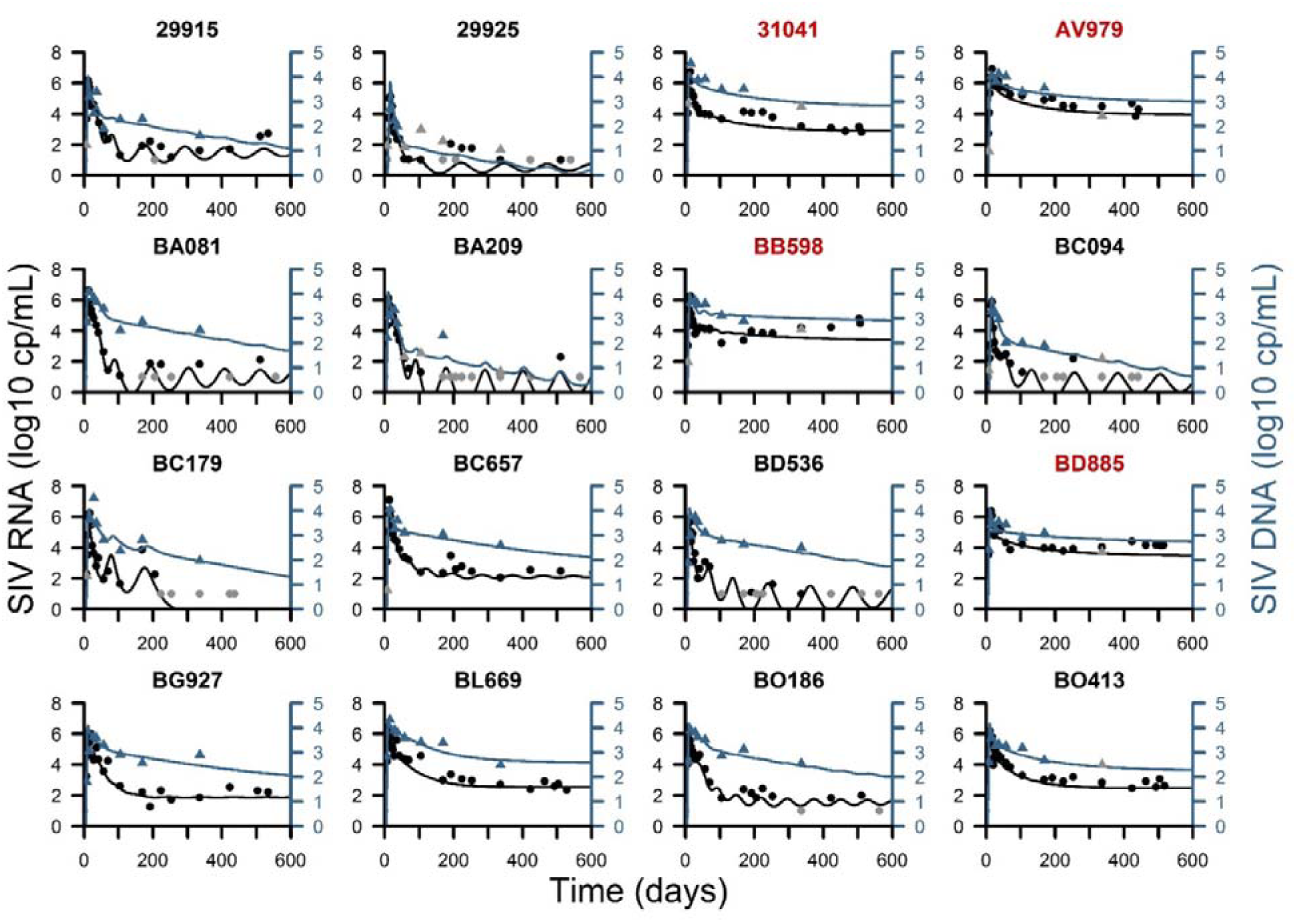
Observations (symbols) and individual fits (solid lines) of SIV-RNA (black, left axis) and SIV-DNA (blue, right axis). Gray symbols are data below the limit of quantification. Black and red legends indicate controllers and viremic macaques, respectively.

The model predicted that the half-life of actively infected cells was equal to 5.5 days and then progressively dropped to 0.3 days upon the establishment of an effective CD8^+^ T-cell cytotoxic response.

The rate of this decline was different between controllers and viremic, with the controllers typically mounting their antiviral response much faster than viremic (Figures 3 and S4). Model predicted that controller reach their maximal level of cytotoxic response within 100 days (median [min – max] time to reach 80% of maximal level was 94 [56 - 167] days), while about 250 days were required in viremic macaques (241 [199 - 364] days). Interestingly this prediction in the change in the actively infected cell half-life matched with the experimental measurements of the *ex vivo* CD8^+^ T-cell antiviral activity in a majority of animals (see Methods and Figure 3), but not with the frequency of SIV-specific CD8^+^ T-cells as determined by intracellular cytokine staining in response to SIV peptides (Figure 4).

**Figure 3:**
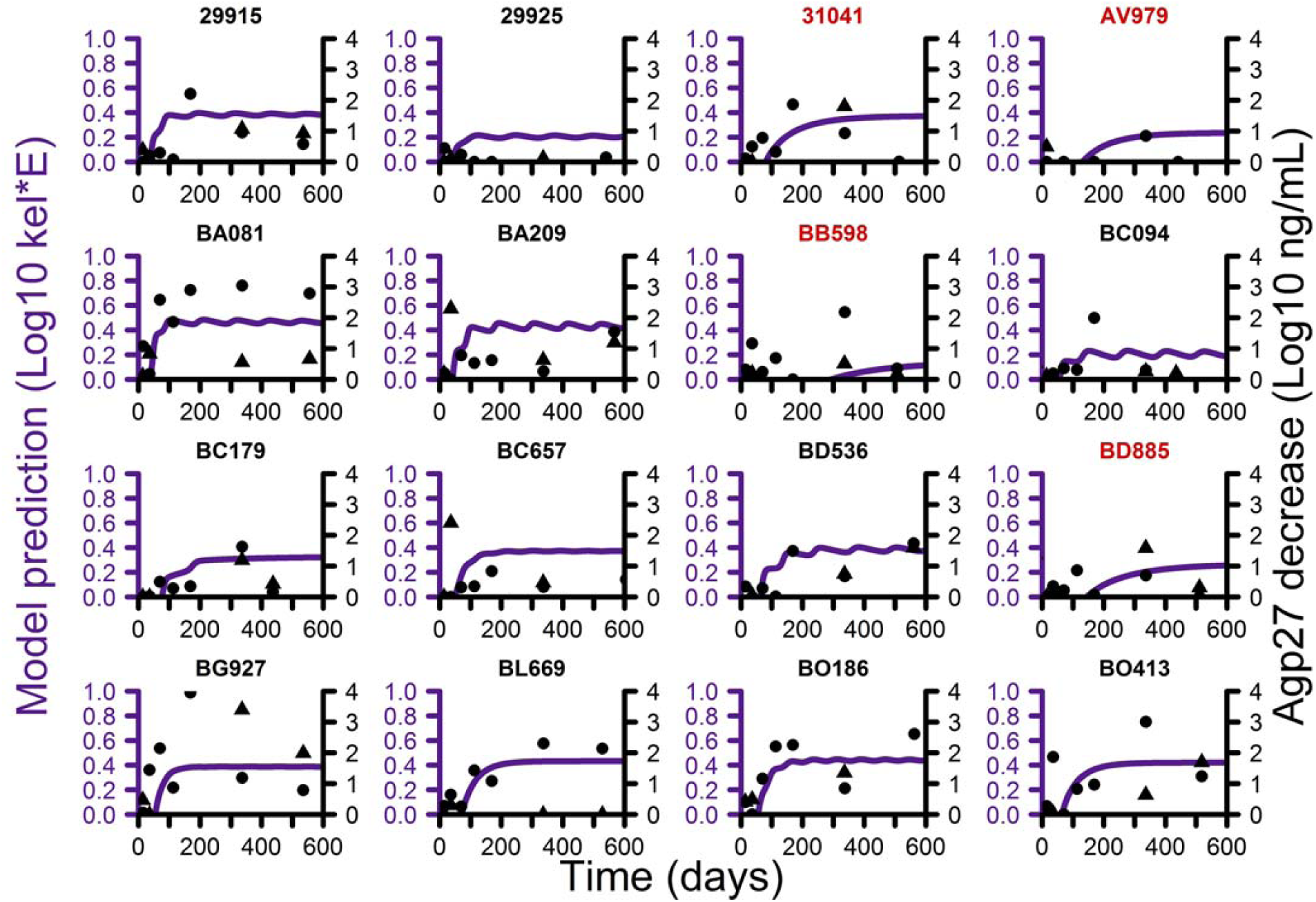
Model individual prediction of the cytotoxic immune response strength (kel*E, purple line, left axis) and observed *ex vivo* CD8^+^ T-cell antiviral activity in PBMC (black dots, right axis) and in peripheral lymph nodes (black triangle, right axis). The identity number of controllers and viremic macaques are denoted in black and red, respectively.

**Figure 4:**
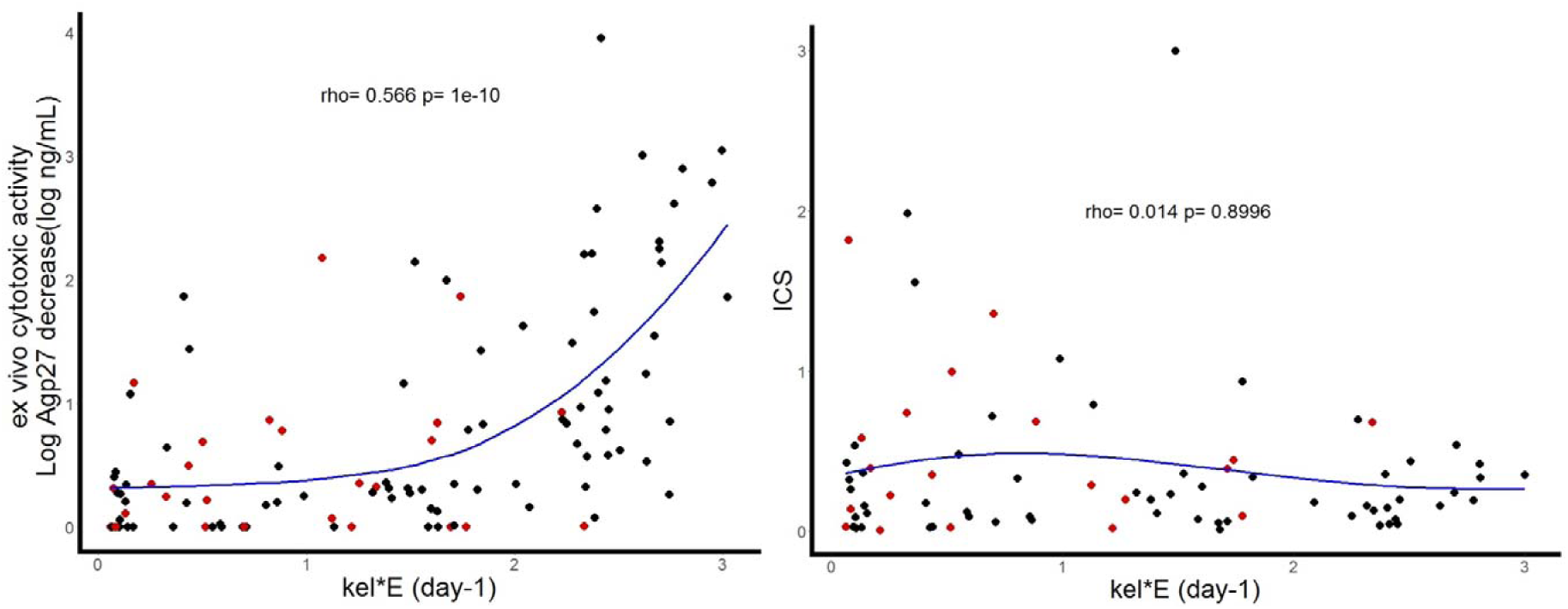
Left: Association between observed *ex vivo* CD8^+^ T-cell antiviral activity and model predictions of cytotoxic immune response strength (kel*E) in all animals. Right: Association between cytokine production measured by intra-cellular staining (ICS). The blue line is the Loess local regression and *ρ* gives the Spearman coefficient regression. Black and red dots are indicative of controller and viremic animals, respectively.

In addition to actively infected cells, the slow decline in SIV-DNA compared to SIV-RNA uncovered two additional compartments of cells that are infected but do not actively produce virus (Figure 5), and we estimated the half-life of these populations to about 5.1 and 118.1 days, respectively (Table 1). We estimated that after encounter and infection, the chance for a target cell to become actively producing, short-lived non-actively producing or long-lived non-actively producing were equal to 47, 52 and 1%, respectively (Table 1). Given the infection rate and the half-life of each compartment, one can also deduce the distribution of these cell populations over time. We estimated that in controllers, about 99.7% of infected cells after day 50 were long-lived non-actively producing cells, only 0.3% are short-lived non-actively producing, and less than 0.1% are actively producing (Figure 5). In contrast, in viremic macaques, whose by definition much higher viral load and more infection events take place, we estimated that after peak viremia about 60% of infected cells are short-lived non-actively producing, 30% are long-lived non-actively producing and about 10% are actively producing cells.

**Table 1:**
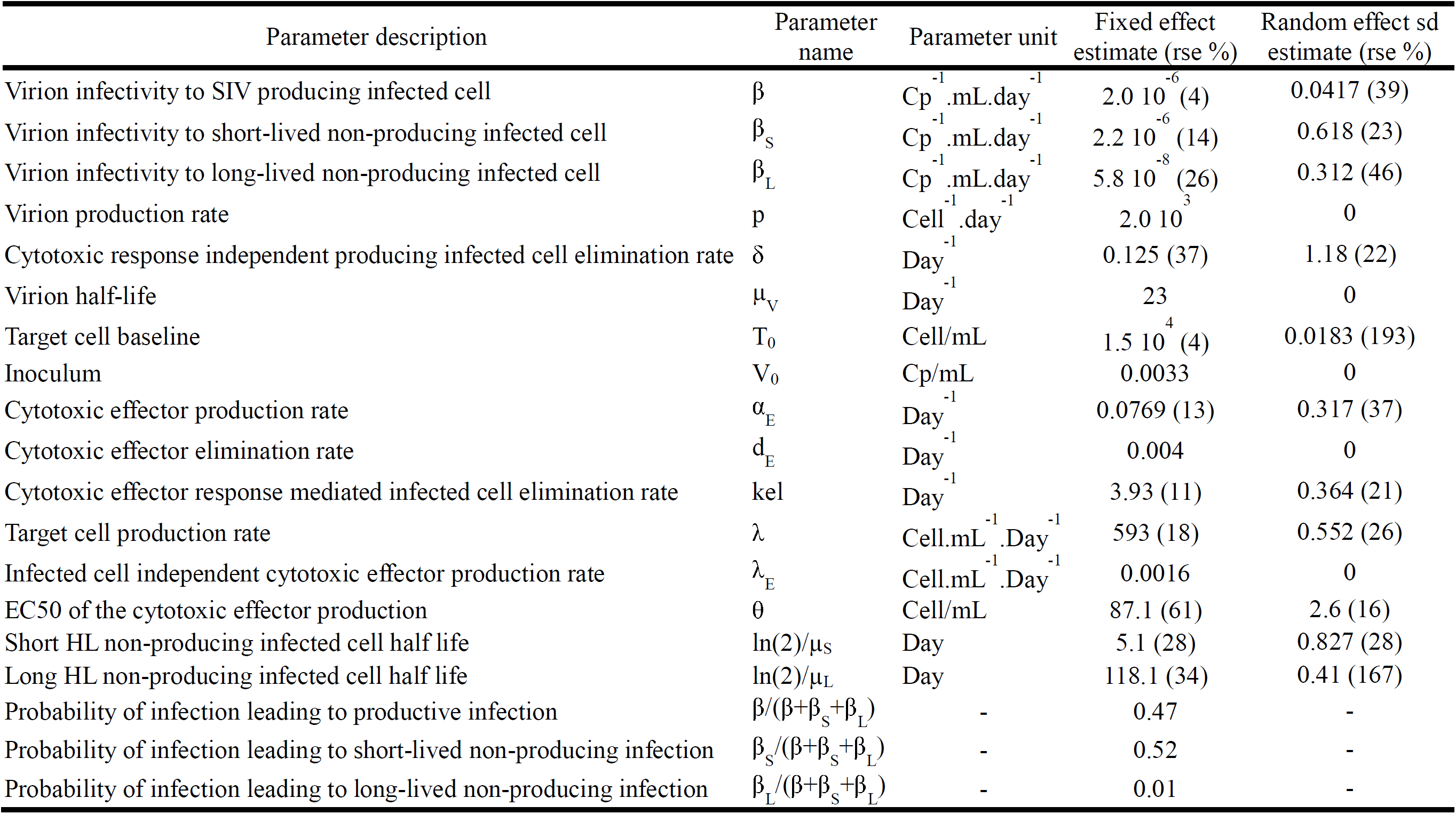
Viral kinetic model parameter estimates (relative standard errors, rse).

**Figure 5:**
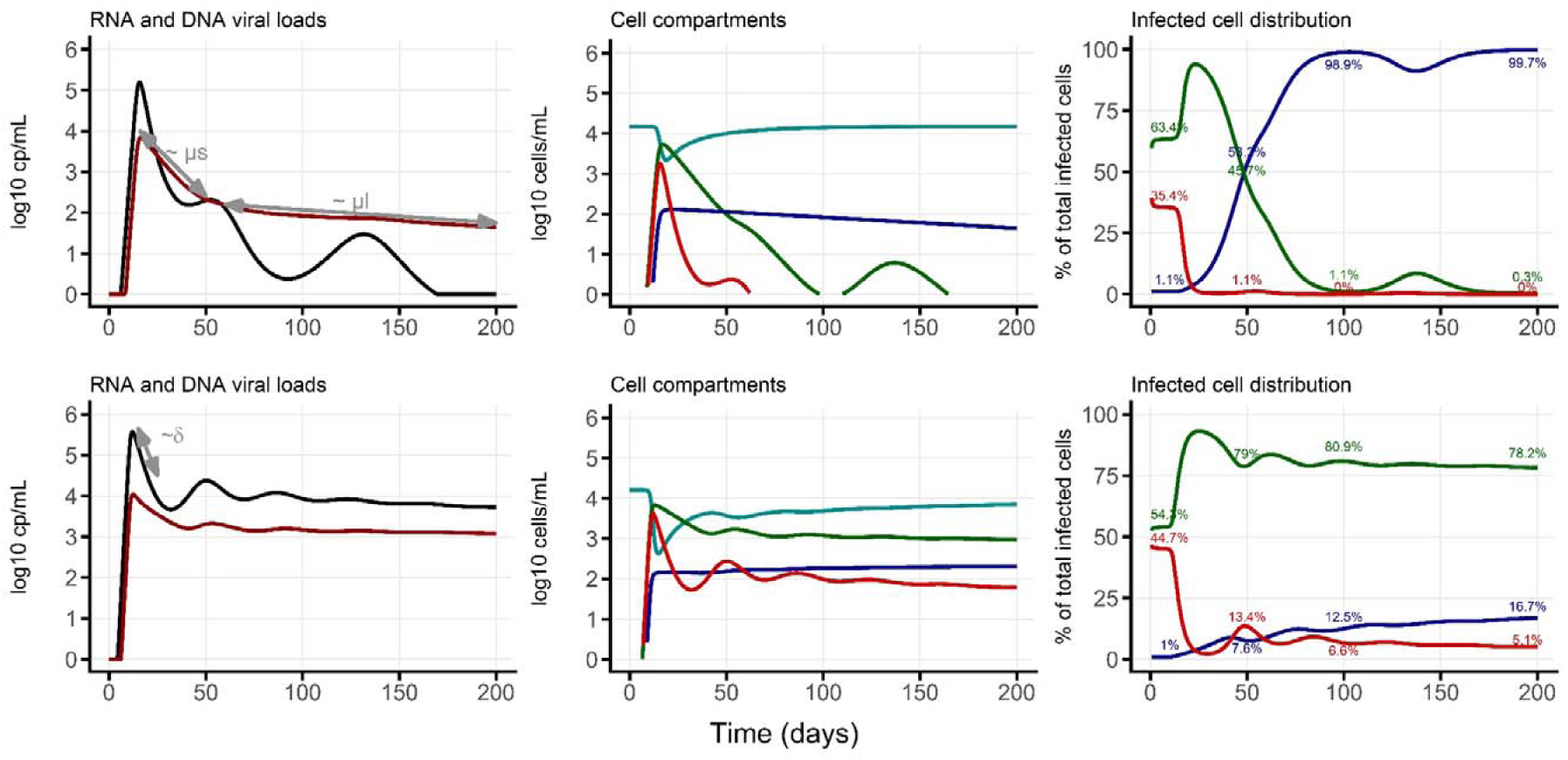
Model predictions in a typical controller (top) or viremic animal (bottom). Left panels: SIV-RNA (black) and SIV-DNA (red) kinetics. Middle panels: cell population kinetics (cyan: target cells, red: actively producing cells, green short-lived non-actively producing cells, blue: long-lived non-actively producing cells). Right panels: Relative contribution to the infected cell population of actively producing (red), short-lived non-actively producing (green) and long-lived non-actively producing (blue). The parameter values used for these simulations correspond to animals BC094 and BB598.

### Exploration of model determinants of viral control

Individual parameter estimates and predicted viral set points were calculated for the 16 macaques, showing a log-linear relationship between θ (EC_50_ of the cytotoxic response development) and the predicted SIV-RNA set point, and illustrating how in this model the viral set point is driven by the kinetics of immune response able to eliminate infected cells (Figure S5). When looking at the property of that model on a larger parameter space, we found that the strength of elimination of infected cells by the immune response, k_el_, was also involved to a lesser extent on the set point value of SIV-RNA (Figures S5 and S6).

## Discussion

In this study, we showed that a mechanistic viral kinetic model incorporating an immune response able to efficiently eliminate infected cells can comprehensively reproduce SIV-RNA and DNA viral kinetics observed in controller and viremic macaques. We predicted that the ability of the immune system to mount such response within three months and reduce the half-life of actively SIV producing cells is the key parameter driving the viral control in this NHP model. Further, by enriching our analysis with the kinetics of SIV-DNA, we found that the best description of the data was provided by a model including three compartments of infected cells or, more exactly, three compartments of SIV-DNA containing cells with distinct half-lives and size. Actively producing infected cells, which are targeted by the immune system had the shortest half-life, with values varying from about 5.8 days initially to 0.3 day after the development of an effective immune response. These cells represented about 50% of the SIV-DNA containing cells initially but rapidly diminish afterwards, consistent with the interpretation that actively producing cells are a small subset of all infected cells. Then we identified a second compartment of cells with a longer half-life of about 5 days, with no statistically significant contribution to SIV-RNA load. This compartment is negligible when SIV-RNA levels are low (<1%) but represents up to 80% of total infected cell population when viremia is large, such as during chronic infection in viremic animals. Lastly, we identified a third compartment with no statistically significant contribution to SIV-RNA load but a long half-life of about 4 months. This compartment represents up to 99.9% of total infected cell population when SIV-RNA levels are low but is negligible when SIV-RNA is large.

Parameter estimates of the model are consistent with those reported in HIV and SIV viral kinetic models. The calculated basic reproductive number, R_0_, which represents the number of new infections resulting from one infected cell, was equal to 10.5 in our study, close to previously reported estimates in HIV or SIV infections with values found between estimate of 8.0 and 13.7 *(24–26)*. Our estimate of initial target cell concentration was 15 cells/µL corresponding to about 2% of the baseline CD4^+^ T-cell counts in blood (median 654 cells/µL). This is consistent with measurement of activated CD4^+^ T-cells, expressing CCR5 at the cell surface (5%) *(27)* or the proliferation marker Ki-67 (1%) *(28)*. In our model the rate of actively infected cell loss in blood is increasing from 0.12 day^−1^ to 3.76 day^−1^ in chronic infection once immune response effectors have reached the steady state, and are in the range of values reported, with 0.39 day^−1^ found in chronic HIV infection *(29)*, and higher values of about 1 day^−1^ in a SIV model of progressors *(30, 31)*. The model relies on the assumption that this increase of loss rate is directly related to the development of the cytotoxic effectors, and what is observed in blood reflects the global process of the infection at the individual scale. The other mechanisms that may lead to loss of infected cells in the peripheral blood compartment, such as necrosis or apoptosis are not explicitly included in the model, and are encompassed in the baseline elimination rate δ.

Although these and other data *(9, 32)* suggested that an efficient cytotoxic immune response is key to achieve viral control, other mechanisms are also likely to contribute to the control viral replication *(9)*, such as neutralizing antibodies or type 1 IFN *(20, 33)*. Further, CD8^+^ T-cell role may not be limited to cytotoxic activity and encompass other functions such as cytokine production *(34–36)*. The model retained here, adapted from *(22)*, assumes that the cytotoxic response is driven by the amount of actively producing infected cells (Figure 1). This indicates that achieving viral control can be possible if a low amount of infected cells are sufficient to trigger an efficient response. If the development of an effective cytotoxic response requires the presence of a large amount of antigen, it may come too late to prevent chronicity. Our hypothesis of a central role of the cytotoxic response is also consistent with measurements of CD8^+^ T-cell cytotoxic activity (Figures 3 and 4). Overall, we found our predictions of the actively producing infected cells half-life to match in several animals the change in *ex vivo* CD8^+^ T-cell dependent direct antiviral activity but not with the frequency of SIV-specific CD8^+^ T cells, indicating that the key determinant of control is not the magnitude *per se* but rather the cytotoxic potential of the CD8^+^ T-cell response *(23)*. However it should be acknowledged that this is not the case for all NHPs. This could be related to insufficient sampling or the role of cytotoxic responses in tissues or other mechanisms involved in the control of viremia that are not reflected in the *ex vivo* cytotoxic activity in blood samples.

The descriptive analysis of the data identified a large discrepancy between SIV-RNA and SIV-DNA kinetics after peak viremia, with SIV-DNA kinetics declining much more slowly than SIV-RNA. Using our model to fit both kinetics simultaneously, we found that the best description of the data was obtained by assuming that SIV-DNA, unlike SIV-RNA, did not only reflect the compartment of actively SIV-producing cells, but also accounted for two additional compartments of infected cells that did not statistically contribute to viral production. What do these two compartments of non-actively producing (more exactly non-producing or low-producing) infected cells represent? The long-lived non-actively producing cell compartment identified in our model has a half-live of about 120 days, visible from the long term SIV-DNA decline observed in controller macaques. This value is close to the lifespan of memory CD4^+^ T-cells *(37)* which is the main population of HIV infected cells in chronic patients or HIV controllers *(18, 38, 39)*, consistent with the idea that long-term viral decline in virally suppressed animals is limited by the long half-life of the latently infected reservoir and their proliferative capacity. A more realistic model incorporating sporadic reactivation processes from this reservoir and new round of infection could not be identified with these data (Figure S7). We estimated that upon encounter and successful infection, the chance to get into this latent compartment is low and close to 1%.

In contrast, the chance to get into short-lived non-actively producing infected compartment were much higher and equal to 52%. The high probability to get into this compartment, together with its short half-life and the absence or low level of viral production suggests that this compartment may correspond to infected cells with an abortive viral cycle (i.e., activated T cells in which infection was not complete and did not lead to virion production called “abortive” hereafter) that enabled pyroptosis *(40)*. Taking advantage of this animal model, which allowed observing a rapid reduction of actively producing cells after peak of viremia, we could precisely estimate the half-life of this compartment to about 5.3 days. Of note, our model only considered infection by free virus and did not incorporate cell to cell infection, which may be another important contributor to abortive infection *(41)* and hence may underestimate the true half-life of this compartment. However it is remarkably consistent with another estimate provided in the context SHIV infection and dynamics in the gut *(42)*, where assumptions on the role of cell to cell infection could be done. Our results also confirm in a SIV pathogenic model the existence of the two slopes decline of viral DNA which was proposed in the non-pathogenic SHIV model *(42)*. Although we found that more than half of cell infection events could be abortive, this could not explain by itself CD4 decline during acute infection. However SIV-DNA do not account for incomplete transcripts *(43)* and thus may only be an underestimate of these “abortively infected cells”. Further we could not distinguish in our data integrated and more labile unintegrated DNA *(44)*, and hence the decay of SIV-DNA kinetics may also account for the loss of unintegrated HIV-DNA. However, current estimates of intracellular HIV-DNA half-life is 2 days, shorter than our estimate of cell loss of 5.3 days and hence should have only a minimal impact on our estimate of the decay rate *(45)*. Another limitation of our model is that we assumed that infected cells had one copy of SIV-DNA. It was suggested that actively producing cells may contain more than one copy of viral DNA per cell *(46)*, whereas non-actively producing or latently infected cells usually stick to one copy per cell *(47)*. A sensitivity analysis performed to address this point demonstrated that for a number of copies per cell ranging from 2 to 20, the selected model was able to provide description of the data as good as with the assumption of 1 copy per cell, but this number of copies per cell cannot be identified from the value of the viral production, ranging from 2,000 to 20,000 cp/cell/day (Table S4). Parameter estimates of the SIV-DNA model remain very close to the ones of the selected model, except a moderate decrease of the short-lived non-actively producing cell elimination rate.

Whether these estimates can be extended to HIV-1 infected individuals is not known, but our data suggest that HIV-DNA kinetics could be a relevant marker of the latent reservoir in individuals with undetectable viremia. However, our results also suggest that as long as viral replication has not been blunted, HIV-DNA represents a mixture of latently and abortively infected cells.

To conclude, we proposed here a mechanistic model to characterize both SIV-RNA and SIV-DNA in viremic and natural controller macaques, where the level of viral set point depends on how fast an efficient cytotoxic immune response is mounted. This model could be used to characterize the decline of infected cell populations during and after antiretroviral therapy to explore the importance of kinetics of various biomarkers in virologic control.

## Materials and Methods

### Study design

Data used in this study were obtained from the ANRS SIC study *(23)*. Briefly, it included 16 male cynomolgus macaques, all infected intrarectally with SIV_mac251_ and followed for 18 months after viral exposure without any treatment. Three experimental groups were initially designed in this study, with predicted increase of control in the long term in two of them *(8, 10)*: i) N=6 without the M6 MHC haplotype and challenged with a high inoculum of 50 AID_50_; ii) N=4 without the M6 haplotype but challenged with a lower inoculum of 5 AID_50_; and iii) N=6 with the M6 haplotype and challenged with a high inoculum of 50 AID_50_.

Consistent with Passaes et al. *(23)*, we defined as “controllers” individuals having two consecutive plasma SIV-RNA lower than 400 cp/mL and with no consecutive SIV-RNA higher than 400 cp/mL afterwards, while others were defined as viremic. With this definition, 12 individuals were defined as controllers (3 in the non-M6/high inoculum, 4 in non-M6/low inoculum, 5 in the M6 groups) and 4 were characterized as viremic (3 in the non-M6/high inoculum, 1 in M6/high inoculum, 0 in the non-H6/low inoculum groups). As described in *(23)*, no differences were found between controllers across groups and thus we considered all controllers as a single group.

### Ethic statement

Mauritius cynomolgus macaques (*Macaca fascicularis*) were housed in the “Commissariat à l’Energie Atomique et aux Energies Alternatives” (CEA, Fontenay-aux-Roses, France) in accordance with French national regulations (CEA Permit Number A 92-032-02). The CEA complies with the Standards for Human Care and Use of Laboratory Animals of the Office for Laboratory Animal Welfare (OLAW, USA) under OLAW Assurance number #A5826-01. All experimental procedures were conducted according to European Directive 2010/63 (recommendation number 9) on the protection of animals used for scientific purposes. This study was accredited under statement number 13-005, by the ethics committee “Comité d’Ethique en Experimentation Animale du CEA” registered under number 44 by the French Ministry of Research. Animals were studied with veterinary guidance, housed in adjoining individual cages allowing social interactions and maintained under controlled conditions with respect to humidity, temperature, and light (12-hour light/12-hour dark cycles). Animals were monitored and fed commercial monkey chow and fruit 1±2 times daily by trained personnel, and water was available ad libitum. Environmental enrichment was provided including toys, novel foodstuffs, and music under the supervision of the CEA Animal Welfare Body. Experimental procedures (animal handling, viral inoculations, samplings) were conducted after sedation with ketamine chlorhydrate (Rhone Merieux, Lyon, France, 10mg/kg). Tissues were collected at necropsy: animals were sedated with ketamine chlorhydrate (Rhone Merieux, Lyon, France, 10mg/kg) then humanly euthanized by intravenous injection of 180mg/kg sodium pentobarbital.

### Data collected

Several virologic and immunologic biomarkers were longitudinally measured to characterize the determinants of control and we only detail below those used in this analysis *(23)*. Plasma SIV-RNA was assessed by real time PCR, with a limit of quantification of 12.3 copies/mL (cp/mL) at days 0, 7, 9, 11, 14, 15, 17, 21, 24, 28 and then weeks 5, 6, 8, 10, 15, 24, 27, 29, 32, 36, 48, 60, 73 post challenge and at euthanasia. SIV-RNA data were converted from cp/mL of plasma to cp/mL of blood assuming a constant hematocrit value of 0.5. Total SIV-DNA was measured in total blood leukocytes using ultrasensitive real time PCR, with a limit of quantification of 2 copies/PCR reaction, at days 0, 7, 15, 28 and weeks 5, 8, 15, 24 and 48 after challenge. This technique allows to quantify all forms of SIV-DNA present in cells, including linear non-integrated DNA, episomal circular DNA, and intra-host genome integrated DNA *(14)*. SIV-DNA levels per million cells were converted from copies/10^6^ leukocytes to cp/mL of blood, using individual blood leukocyte counts sampled simultaneously to the SIV-DNA measurements. The frequency of SIV-specific CD8^+^ T cells was determined by intracellular cytokine staining in response to SIV peptides. A qualitative measure of the direct antiviral efficacy of the SIV-specific CD8^+^ T-cell immune response was assessed following a procedure detailed in *(48)* and adapted to the cynomolgus macaque/SIV model *(8)*. Briefly, it consists in the difference in p27 antigen production by CD4^+^ T-cells infected with SIVmac_251_ in absence and in presence of autologous CD8^+^ T-cells obtained from infected macaque tissue. This *ex-vivo* cytotoxic activity was measured in blood 3 weeks before challenge and then 2, 5, 10, 16, 24, 48 weeks after challenge and at euthanasia, and in lymphatic nodes and spleen with a sparse sampling protocol.

### Exploratory analysis

Peak value, time to peak, slope between the peak and day 168 post-infection and mean value after day 168 post-infection were calculated for SIV-RNA, SIV-DNA, SIV-RNA/DNA ratio and *ex-vivo* CD8^+^ T-cell activity. All metrics were compared between controllers and viremic animals using non-parametric Wilcoxon test without correction for test multiplicity.

### Strategy for model selection of the cytotoxic immune response

Based on previous results obtained in SIV and HIV *(1, 3, 7, 9)*, we considered an adaptation of the target cell limited model accounting for the cellular cytotoxic immune response. In order to find the best parameterization of the model, we identified 9 candidate models from the literature (Table S2) *(22, 33, 49–51)*. For the sake of parameter identifiability we fixed in all models the free virion elimination rate, µ_V_, to 23 day^−1^ *(52)* and the viral production rate per cell, p, to 2,000 day^−1^ *(22)*. Because some models contain parameters that are poorly identifiable, we used a three steps procedure to determine the best possible fitting to the data that can be achieved with each model. First, we generated 200 sets of initial parameters using Sobol pseudo random sequences *(53)*, taking as boundaries of sampling intervals the reported value of the parameters of interest +/- 2 log10. Second, estimation was performed for each set of initial parameters and the 5 estimations with the lowest Bayesian Information Criterion (BIC, the lower the better) were retained and used as initial parameters to provide the final set of parameter estimates. When parameters were not identifiable or associated with a relative standard error larger than 50%, they were fixed to the value found at the first step. Finally, the lowest BIC obtained for each candidate model was retained and the model providing the lowest BIC over all candidate models was retained and carried forward. All model estimations were performed using SAEM algorithm implemented in Monolix software (http://lixoft.com/).

### Extension of the model to SIV-DNA dynamics

Following conclusion of a previous report *(47)*, we assumed that one DNA copy corresponds to one infected cell. The model selected at the previous stage was first used to fit SIV-RNA and SIV-DNA kinetics simultaneously, assuming that SIV-DNA reflects the number of actively infected cells, as previously proposed *(42)*. Because this model could not adequately fit SIV-DNA kinetics (see results), we extended it assuming that the total number of infected cells also encompassed i) non-actively HIV producing cells with or without cell division and/or; ii) non-actively producing cells that reactivate into actively producing cells after a latency period. Model selection was based on BIC.

### Exploration of model determinants of viral control

In order to evaluate the parameters that were associated with viral control, Spearman correlation coefficient was computed between individual parameters and viral set point (mean SIV-RNA observed after day 168 post-infection). Then we explored the associations in a larger parameter space using a Sobol global sensitivity analysis *(54)* and a range of sampled value defined as the population value +/- one standard deviation of the random effect. The R package sensitivity was used for the Sobol analysis (https://cran.r-project.org/web/packages/sensitivity/index.html).

### Association of model prediction to *ex vivo* SIV-specific CD8^+^ T cell cytotoxic activity

For each macaque, we calculated the cytotoxic response mediated infected cell elimination rate predicted by the model, and given by E(t) x k_el_ (see results). Model predictions and *ex vivo* CD8 antiviral activity measured in peripheral blood mononuclear cells (PBMC) and lymphatic nodes were plotted over time, assuming an arbitrary 1:1 ratio. Then, the association between model predictions and *ex vivo* CD8 antiviral activity in PBMC, and between model predictions and frequency of SIV-specific CD8^+^ T-cells, at each sample time, was assessed using Spearman correlation test.

## List of Supplementary Materials

Table S1: SIV-RNA, SIV-DNA, SIV-RNA/DNA ratios and *ex vivo* CD8^+^ T-cell antiviral activity in cynomolgus macaques infected with SIV_mac251_

Table S2: List of the different models tested to fit SIV-RNA kinetics, reference in the literature, and best fitting criterion obtained

Table S3: Model used to fit SIV-RNA and SIV-DNA kinetics and associated BIC.

Table S4: Sensitivity analysis evaluating the variation of the number of SIV-DNA copies per actively infected cell.

Figure S1: Plasma SIV-RNA, SIV-DNA in blood, *ex vivo* CD8^+^ T-cell antiviral activity and SIV-RNA/DNA ratio over time in controllers and viremic macaques

Figure S2: Individual fits of SIV-DNA kinetics assuming that all infected cells are actively producing, one single compartment of non-actively producing cells and two compartments of non-actively producing cells.

Figure S3: Observation and individual fit obtained by individual regression of SIV-DNA kinetics after peak viremia using a mono-exponential function or a bi-exponential function.

Figure S4: Model individual prediction of cytotoxic immune response strength (kel*E) over time.

Figure S5: Association between individual values of model parameters obtained by empirical Bayesian estimates and individual predicted viral set point.

Figure S6: First order and total order Sobol sensitivity indices on viral set point for SIV-RNA model parameter.

Figure S7: Model prediction assuming a reactivation rate of long-lived non-actively producing cells.

## Acknowledgements

We thank Alan Perelson, Ruy Ribeiro and Rodolphe Thiébaut for their helpful comments and suggestions on this work.

