## supplementary table and figures for "Modeling SIV kinetics supports that cytotoxic response drives natural control and unravels heterogeneous populations of infected cells"

**One sentence summary:** Modeling viral dynamics in SIV natural controller macaques predicts that viral control is primarily driven by the capability to establish an efficient cytotoxic response and the viral decline during control unravels distinct compartments of infected cells.

**Authors:**

V. Madelain^1^, C. Passaes^2,3^, A. Millet^4^, V. Avettand-Fenoel^4^, R. Djidjou-Demasse^5^, N. Dereuddre-Bosquet^3^, R. Le Grand^3^, C. Rouzioux^4^, B. Vaslin^3^, A. Saez-Cirion^2^, J. Guedj^1^

**Affiliations:**

^1^IAME, UMR 1137, INSERM, Université Paris Diderot, Sorbonne Paris Cité Paris, France

^2^Institut Pasteur, Unité HIV, Inflammation et Persistance, Paris, France

^3^CEA, Université Paris Sud, INSERM U1184, Immunology of Viral Infections and Autoimmune Diseases (IMVA), IDMIT Department / IBFJ, 92265 Fontenay-aux-Roses, France.

^4^Université Paris-Descartes, Sorbonne Paris-Cité, Faculté de Médecine, EA 7327, Paris, France

^5^UMR 1065, INRA, Villenave d’Ornon F-33882, France.

List of Supplementary Materials:

Table S1: SIV-RNA, SIV-DNA, SIV-RNA/DNA ratios and *ex vivo* CD8^+^ T-cell antiviral activity in cynomolgus macaques infected with SIV_mac251_

Table S2: List of the different models tested to fit SIV-RNA kinetics, reference in the literature, and best fitting criterion obtained

Table S3: Model used to fit SIV-RNA and SIV-DNA kinetics and associated BIC.

Table S4: Sensitivity analysis evaluating the variation of the number of SIV-DNA copies per actively infected cell.

Figure S1: Plasma SIV-RNA, SIV-DNA in blood, *ex vivo* CD8^+^ T-cell antiviral activity and SIV-RNA/DNA ratio over time in controllers and viremic macaques

Figure S2: Individual fits of SIV-DNA kinetics assuming that all infected cells are actively producing, one single compartment of non-actively producing cells and two compartments of non-actively producing cells.

Figure S3: Observation and individual fit obtained by individual regression of SIV-DNA kinetics after peak viremia using a mono-exponential function or a bi-exponential function.

Figure S4: Model individual prediction of cytotoxic immune response strength (kel*E) over time.

Figure S5: Association between individual values of model parameters obtained by empirical Bayesian estimates and individual predicted viral set point.

Figure S6: First order and total order Sobol sensitivity indices on viral set point for SIV-RNA model parameter.

Figure S7: Model prediction assuming a reactivation rate of long-lived non-actively producing cells.

**Supplementary material:**

**Table S1:** SIV-RNA, SIV-DNA, SIV-RNA/DNA ratios and *ex vivo* CD8^+^ T-cell antiviral activity in cynomolgus macaques infected with SIVmac251.

|  | **All macaques** | | **Controller** | | **Viremic** | |  |
| --- | --- | --- | --- | --- | --- | --- | --- |
| **SIV-RNA** | median | *[min;max]* | median | *[min;max]* | median | *[min;max]* | p |
| **Peak (log cp/mL)** | 6.20 | *[5.13; 7.09]* | 6.14 | *[5.13; 7.09]* | 6.40 | *[6.24; 6.92]* | 0.058 |
| **time to peak (days)** | 14 | *[11; 17]* | 12.5 | *[11; 17]* | 14 | *[14; 17]* | 0.27 |
| **Descending slope (log cp/mL/day)** | -0.038 | *[-0.061; -0.015]* | -0.036 | *[-0.061; -0.018]* | -0.013 | *[-0.028; -0.015]* | **0.013** |
| **Mean viral load after D168 (log cp/mL)** | 1.90 | *[0.26; 4.53]* | 1.25 | *[0.26; 2.88]* | 3.80 | *[2.89; 4.53]* | **0.0011** |
| **SIV-DNA** |  |  |  |  |  |  |  |
| **Peak (log cp/mL)** | 3.73 | *[2.7;4.6]* | 3.7 | *[2.7;4.5]* | 3.9 | *[3.5;4.6]* | 0.38 |
| **time to peak (days)** | 21.5 | *[15;36]* | 21.5 | *[15;36]* | 32 | *[15;36]* | 0.15 |
| **Descending slope (log cp/mL/day)** | -0.011 | *[-0.023;-0.0032]* | -0.0040 | *[-0.023;-0.0032]* | -0.0041 | *[-0.011;-0.0071]* | 0.17 |
| **Mean viral load after D168 (log cp/mL)** | 2.7 | *[1.22;3.16]* | 2.49 | *[1.22; 2.93]* | 2.83 | *[2.71; 3.16]* | **0.042** |
| **SIV-RNA/DNA** |  |  |  |  |  |  |  |
| **Peak** | 2.0 | *[1.1; 2.4]* | 2.0 | *[1.1;2.35]* | 2.0 | *[1.28;2.38]* | 0.77 |
| **time to peak (days)** | 15 | *[7.36]* | 15 | *[7;36]* | 15 | *[15;15]* | 0.58 |
| **Descending slope (/day)** | -0.0133 | *[-0.022; 0.00049]* | -0.015 | *[-0.022;-0.00049]* | -0.0054 | *[-0.0087;-0.0037]* | **0.013** |
| ***Ex vivo* activity** |  |  |  |  |  |  |  |
| **Peak (log ng/mL)** | 1.93 | *[0.44; 3.95]* | 2.10 | *[0.44;3.95]* | 1.36 | *[0.84-2.18]* | 0.32 |
| **time to peak (days)** | 337 | *[15; 567]* | 337 | *[15;567]* | 253 | *[113;337]* | 0.49 |
| **ascending slope (log ng/mL/day)** | 0.0010 | *[-0.00052;0.0039]* | 0.00176 | *[-0.00052;0.0039]* | 0.00038 | *[-0.00032;0.0008]* | 0.13 |

Controller group was defined with SIV-RNA below 400 cp/mL at day 168 post-infection and no consecutive viral loads above afterwards, others were defined as viremic. For SIV-RNA and SIV-DNA, data were log10 transformed, viral AUC was calculated from day 0 to day 511 post-infection, and descending slope were computed from time to peak to day 168 post-infection , as well as for SIV-RNA/DNA ratio. *Ex vivo* CD8^+^ T-cell antiviral activity was computed from day 0 to last sampling. Groups were compared using Wilcoxon test.

**Table S2:** List of the different models tested to fit SIV-RNA kinetics, reference in the literature, and best fitting criterion obtained (see Material and Methods section).

| **Model** | **Reference** | **Use** | **Virus** | $\frac{\boldsymbol{dE}}{\boldsymbol{dt}}$ | $\frac{\boldsymbol{dI}}{\boldsymbol{dt}}$ | **Objective function** | **BIC** |
| --- | --- | --- | --- | --- | --- | --- | --- |
| 0 | *De Boer and Perelson JTB 1998* | reproduced | HIV | _ | $\beta TV-\delta I$ | 944.7 | 969.7 |
| 1 | *Burg et al JTB 2009* | reproduced | HIV | $\frac{\alpha_{E}I}{(I+\theta)}-d_{E}E$ | $\beta TV-\delta I-k_{el}EI$ | 827.4 | 869.0 |
| 2 | *Bonhoeffer et al AIDS 2000* | reproduced | HIV | $\alpha_{E}I-d_{E}E$ | $\beta TV-\delta I-k_{el}EI$ | 845.0 | 881.1 |
| 3 | *Bonhoeffer et al AIDS 2000* | reproduced | HIV | $\frac{\alpha_{E}IE}{(I+\theta)}-d_{E}E$ | $\beta TV-\delta I-k_{el}EI$ | 830.3 | 871.9 |
| 4 | *Li and Handel JTB 2014* | reproduced | influenza | $\lambda_{E}+\alpha_{E}I(E_{max}-E)-d_{E}E$ | $\beta TV-\delta I-k_{el}EI$ | 853.2 | 894.8 |
| 5 | *Li and Handel JTB 2014* | adapted | influenza | $\frac{\alpha_{E}I}{(I+\theta)}-d_{E}E$ | $\beta TV-\delta I-\frac{k_{el}EI}{(E+\zeta)}$ | 805.1 | 852.2 |
| 6 | *Conway and Perelson PNAS 2015* | adapted | HIV | $\lambda_{E}+\alpha_{E}E-d_{E}E$ | $\beta TV-\delta I-k_{el}EI$ | 871.7 | 907.7 |
| 7 | *Conway and Perelson PNAS 2015* | adapted | HIV | $\lambda_{E}+ \frac{\alpha_{E}EI}{(I+\theta)}-d_{E}E$ | $\beta TV-\delta I-k_{el}EI$ | 804.2 | 845.8 |
| 8 | *Conway and Perelson PNAS 2015* | adapted | HIV | $\lambda_{E}- \frac{\delta_{E}EI}{(I+\theta)}-d_{E}E$ | $\beta TV-\delta I-k_{el}EI$ | 870.0 | 911.6 |
| 9 | *Conway and Perelson PNAS 2015* | reproduced | HIV | $\lambda_{E}+ \frac{\alpha_{E}EI}{(I+\theta1)}- \frac{\delta_{E}EI}{(I+\theta2)}-d_{E}E$ | $\beta TV-\delta I-k_{el}EI$ | 797.7* | 844.9* |

*non-identifiable.

**Table S3:** Model used to fit SIV-RNA and SIV-DNA kinetics and associated BIC.

| **Model** | **Formula**  $DNA VL={log}_{10}(I+L+S)$ | **Description** | **BIC** | **LRT**  **p value** | **Residual**  **SIV-RNA**  **(log10 cp/mL)** | **Residual**  **SIV-DNA**  **(log10 cp/mL)** |
| --- | --- | --- | --- | --- | --- | --- |
| 1 | $\frac{dI}{dt}=\beta TV-\delta I-k_{el}EI$ | All infected cells I are productive | 1133.6 | _ | 0.460 | 1.68 |
| 2 | $\frac{dI}{dt}=\beta TV-\delta I-k_{el}EI$  $\frac{dL}{dt}=\beta_{L}TV-\mu_{L}L$ | Addition of a non-productively infected cell compartment L, not targeted by immune response E | 923.4 | <10^-5^§ | 0.481 | 0.467 |
| 2a | $\frac{dI}{dt}=\beta TV-\delta I-k_{el}EI$  $\frac{dL}{dt}=\beta_{L}TV-\mu_{L}L+\rho L$ | Addition of a non-productively infected cell compartment L, able to proliferate, not targeted by immune response E | 931.8 | 1* | 0.478 | 0.472 |
| 2b | $\frac{dI}{dt}=\beta TV-\delta I-k_{el}EI+aL$  $\frac{dL}{dt}=\beta_{L}TV-\mu_{L}L-aL$ | Addition of a non-productively infected cell compartment L, able to reactivate into I, not targeted by immune response E | 919.2 | 0.007* | 0.486 | 0.479 |
| 3 | $\frac{dI}{dt}=\beta TV-\delta I-k_{el}EI$  $\frac{dL}{dt}=\beta_{L}TV-\mu_{L}L$  $\frac{dS}{dt}=\beta_{S}TV-\mu_{S}S$ | Addition of two non-productively infected cell compartments L and S, with long and short half-lives respectively, not targeted by immune response E | **913.5** | **3.0 10^-5^*** | **0.463** | **0.397** |
| § LRT vs model 1, * LRT vs model 2  The parameter *a* is the transfer rate from non-actively infected to actively infected compartment, and ρ the proliferation rate. | | | | | | |

**Table S4:** Sensitivity analysis evaluating the variation of the number of SIV-DNA copies per actively infected cell.

| Viral production (cp/cell/day) | DNA cp per actively productive cell | DNA cp per short-lived non-productive cell | DNA cp per long-lived non-productive cell | OF | BIC | Residual error SIV-RNA | Residual error SIV-DNA | Infectivity constant of long-lived non-productive cell | Infectivity constant of short-lived non-productive cell | Elimination rate of long-lived non-productive cell (/day) | Elimination rate of short-lived non-productive cell (/day) |
| --- | --- | --- | --- | --- | --- | --- | --- | --- | --- | --- | --- |
| 2000 | 1 | 1 | 1 | 847 | 913.5 | 0.463 | 0.397 | -7.24 | -5.66 | 0.0059 | 0.136 |
| *2000* | *5* | *1* | *1* | *855.6* | *922.1* | *0.47* | *0.455* | *-6.92* | *-6.53* | *0.01* | *NE* |
| **4000** | **5** | **1** | **1** | **842.3** | **908.9** | **0.469** | **0.411** | **-6.6** | **-5.8** | **0.0092** | **0.5** |
| 8000 | 5 | 1 | 1 | 842.7 | 909.3 | 0.467 | 0.401 | -6.67 | -5.51 | 0.0076 | 0.09 |
| 10000 | 5 | 1 | 1 | 846.1 | 912.6 | 0.469 | 0.409 | -6.91 | -5.53 | 0.0043 | 0.045 |
| 12000 | 5 | 1 | 1 | 851.5 | 918.1 | 0.467 | 0.388 | -6.74 | -5.38 | 0.0059 | 0.078 |
| 20000 | 5 | 1 | 1 | 855.4 | 921.9 | 0.469 | 0.416 | -6.88 | -5.35 | 0.0041 | 0.039 |
| *2000* | *10* | *1* | *1* | *885.3* | *951.8* | *0.468* | *0.515* | *-6.87* | *-5.74* | *0.0081* | *NE* |
| *4000* | *10* | *1* | *1* | *852* | *918.5* | *0.47* | *0.445* | *-6.51* | *-5.84* | *0.01* | *NE* |
| 8000 | 10 | 1 | 1 | 845.5 | 912.1 | 0.462 | 0.412 | -7.14 | -5.61 | 0.0029 | 0.049 |
| **10000** | **10** | **1** | **1** | **844.4** | **910.9** | **0.463** | **0.412** | **-6.84** | **-5.63** | **0.0055** | **0.05** |
| 12000 | 10 | 1 | 1 | 845.4 | 912 | 0.463 | 0.408 | -6.35 | -5.61 | 0.009 | 0.07 |
| 20000 | 10 | 1 | 1 | 855.3 | 921.8 | 0.466 | 0.434 | -6.66 | -5.48 | 0.0064 | 0.03 |
| *2000* | *20* | *1* | *1* | *919.9* | *986.4* | *0.481* | *0.629* | *-6.89* | *-6.64* | *0.0066* | *NE* |
| *4000* | *20* | *1* | *1* | *880.7* | *947.3* | *0.477* | *0.511* | *-6.71* | *-5.75* | *0.0067* | *NE* |
| *8000* | *20* | *1* | *1* | *857.6* | *924.1* | *0.469* | *0.442* | *-6.29* | *-5.72* | *0.00871* | *NE* |
| 10000 | 20 | 1 | 1 | 847.9 | 914.5 | 0.46 | 0.436 | -6.68 | -5.74 | 0.0074 | 0.08 |
| 12000 | 20 | 1 | 1 | 848.8 | 915.3 | 0.472 | 0.422 | -6.89 | -5.62 | 0.0048 | 0.035 |
| **20000** | **20** | **1** | **1** | **844** | **910.6** | **0.464** | **0.41** | **-6.58** | **-5.58** | **0.0075** | **0.047** |

OF: objective function, BIC Bayesian information criterion, NE could not be estimated.

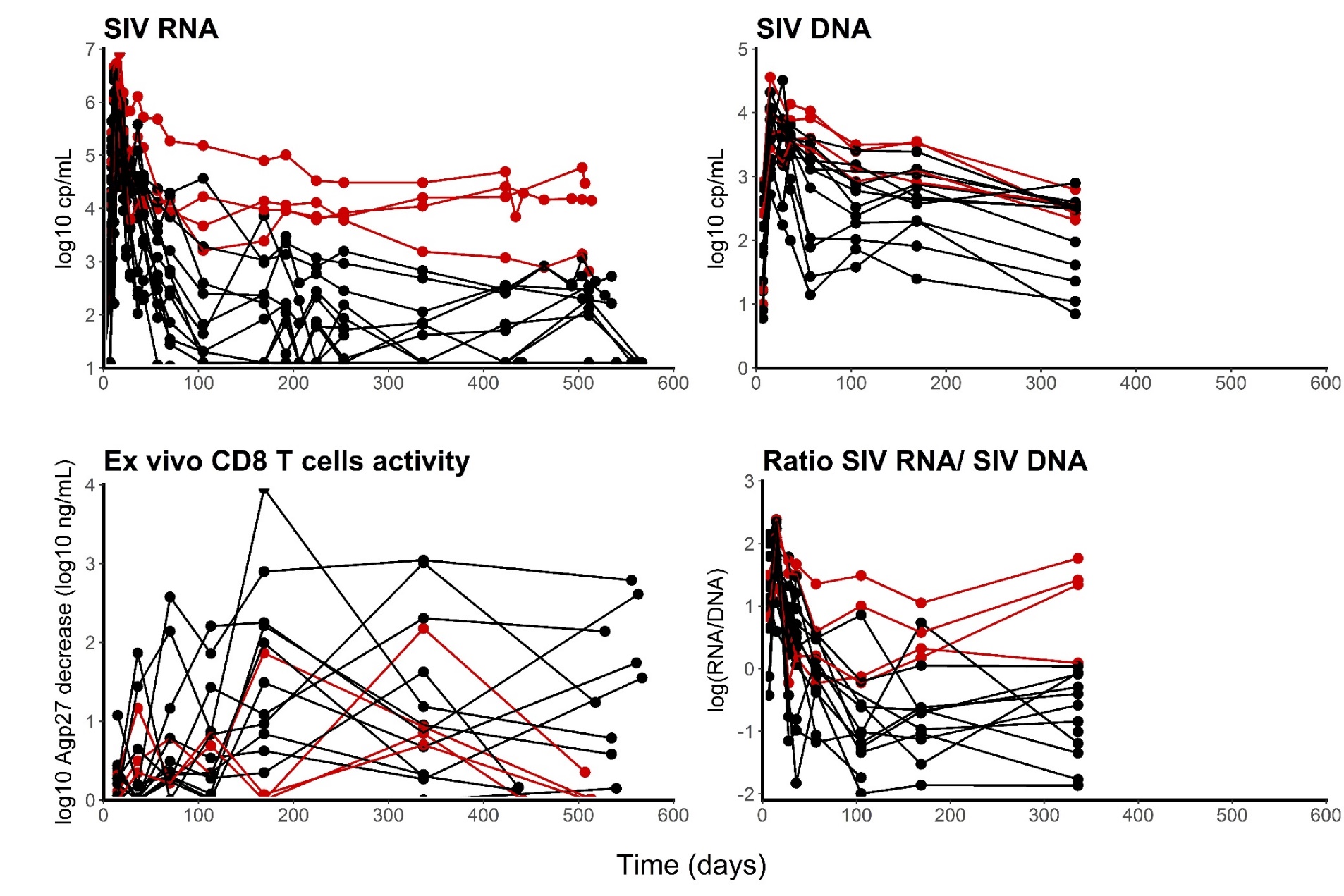

**Figure S1:** Plasma SIV-RNA (top left), SIV-DNA in blood (top right), *ex vivo* CD8^+^ T-cell antiviral activity (bottom left) and SIV-RNA/DNA ratio (bottom right) over time in controllers (black) and viremic macaques (red) as described in *(23)*.

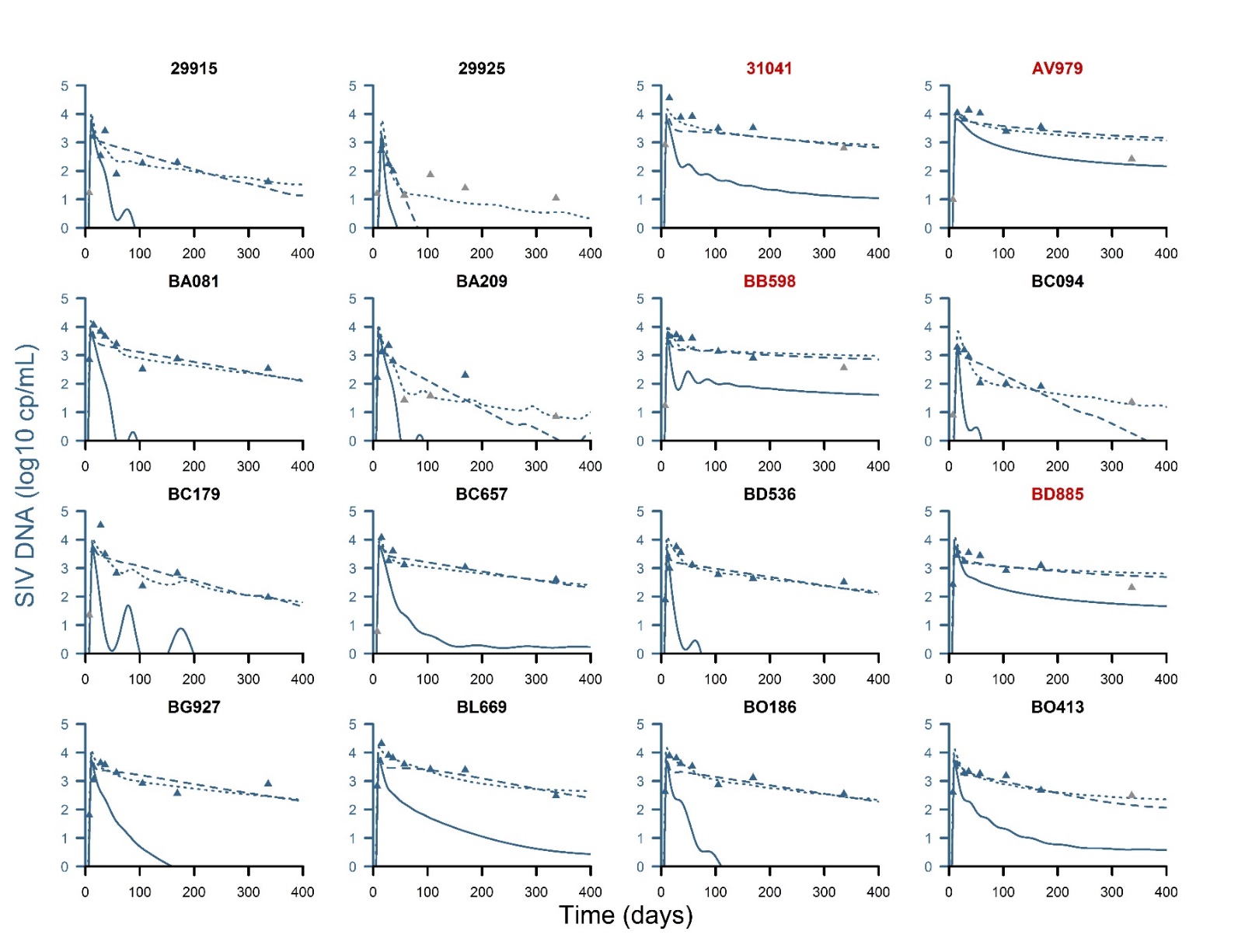

**Figure S2:** Individual fits of SIV-DNA kinetics assuming that all infected cells are actively producing (solid line), and assuming only a single compartment of non-actively producing cells (dashed lines). The best model (see results) is obtained assuming two compartments of non-actively producing cells (dotted lines). Gray symbols correspond to data below the limit of quantification. Black and red legends indicate controllers and viremic macaques, respectively.

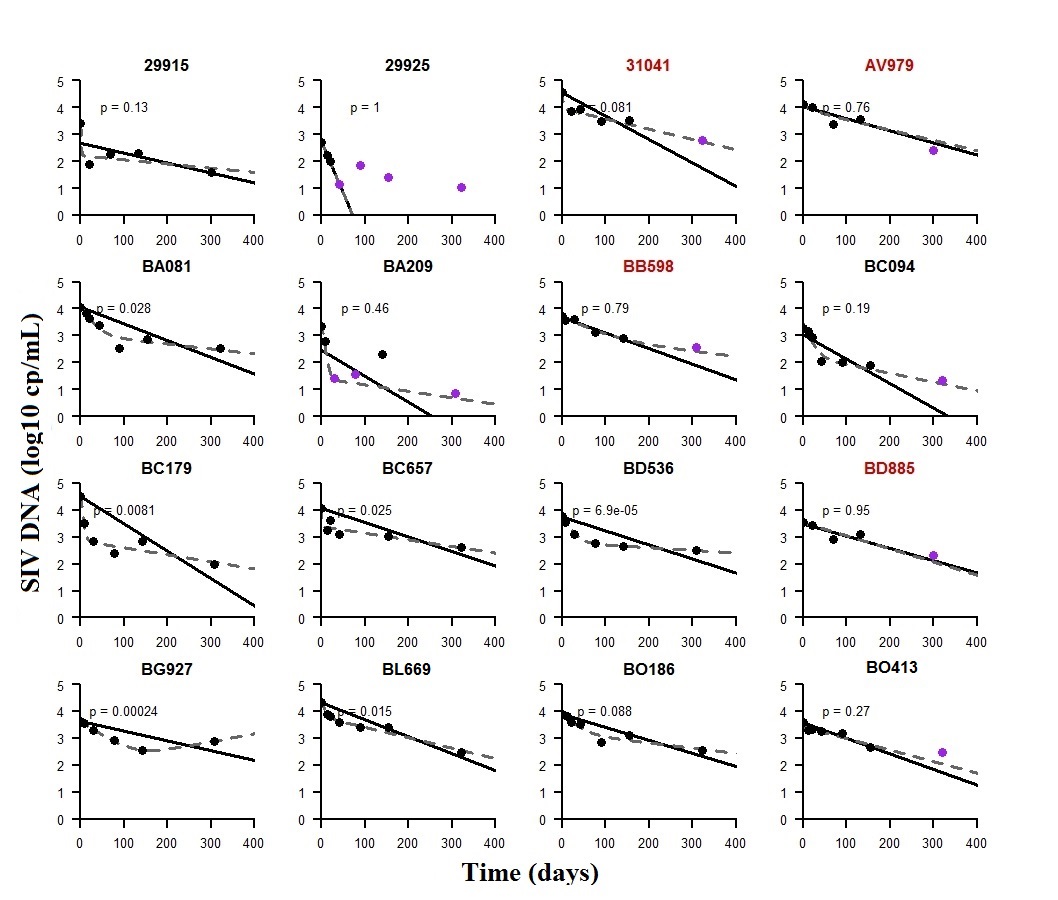

**Figure S3:** Observation (dots) and individual fit obtained by individual regression of SIV-DNA kinetics after peak viremia using a mono-exponential function (solid line) or a bi-exponential function (dashed line). Purple correspond to data below the limit of quantification. P-value <0.05 indicate that the bi-exponential function provides a statistically better fit to the data than the mono-exponential function (likelihood ratio test). Black and red legends indicate controllers and viremic macaques, respectively.

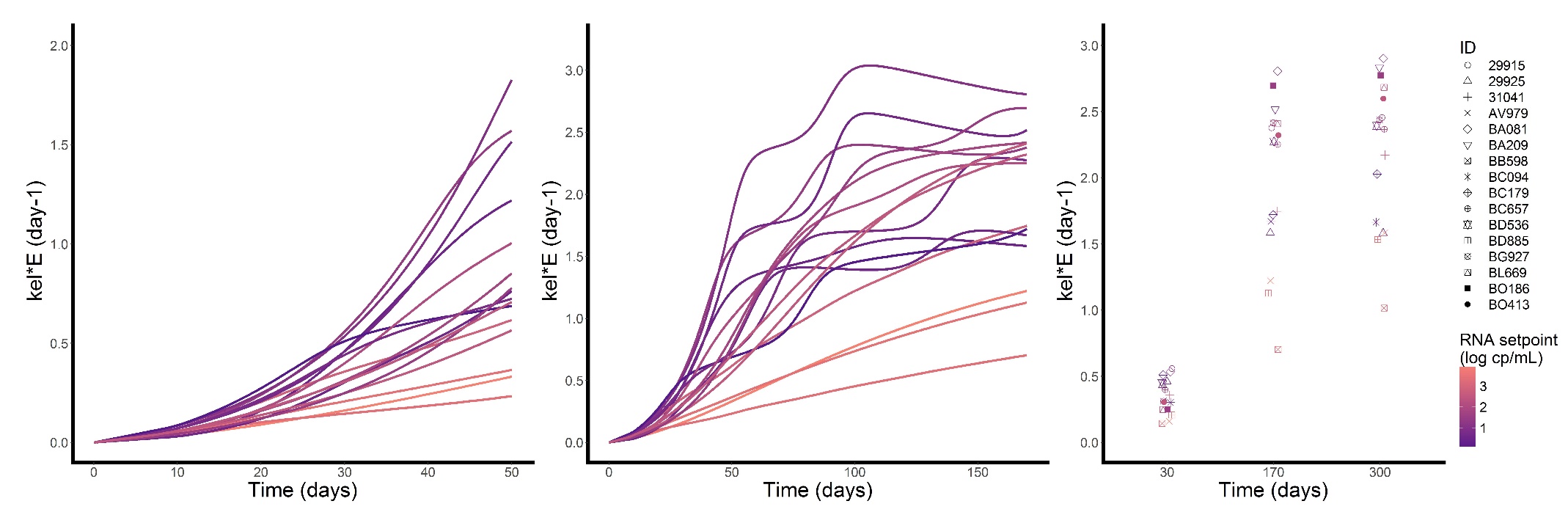

**Figure S4:** Model individual prediction of cytotoxic immune response strength (kel*E) over time. The darker is the line color, the lower is the predicted viral set point of the macaque.

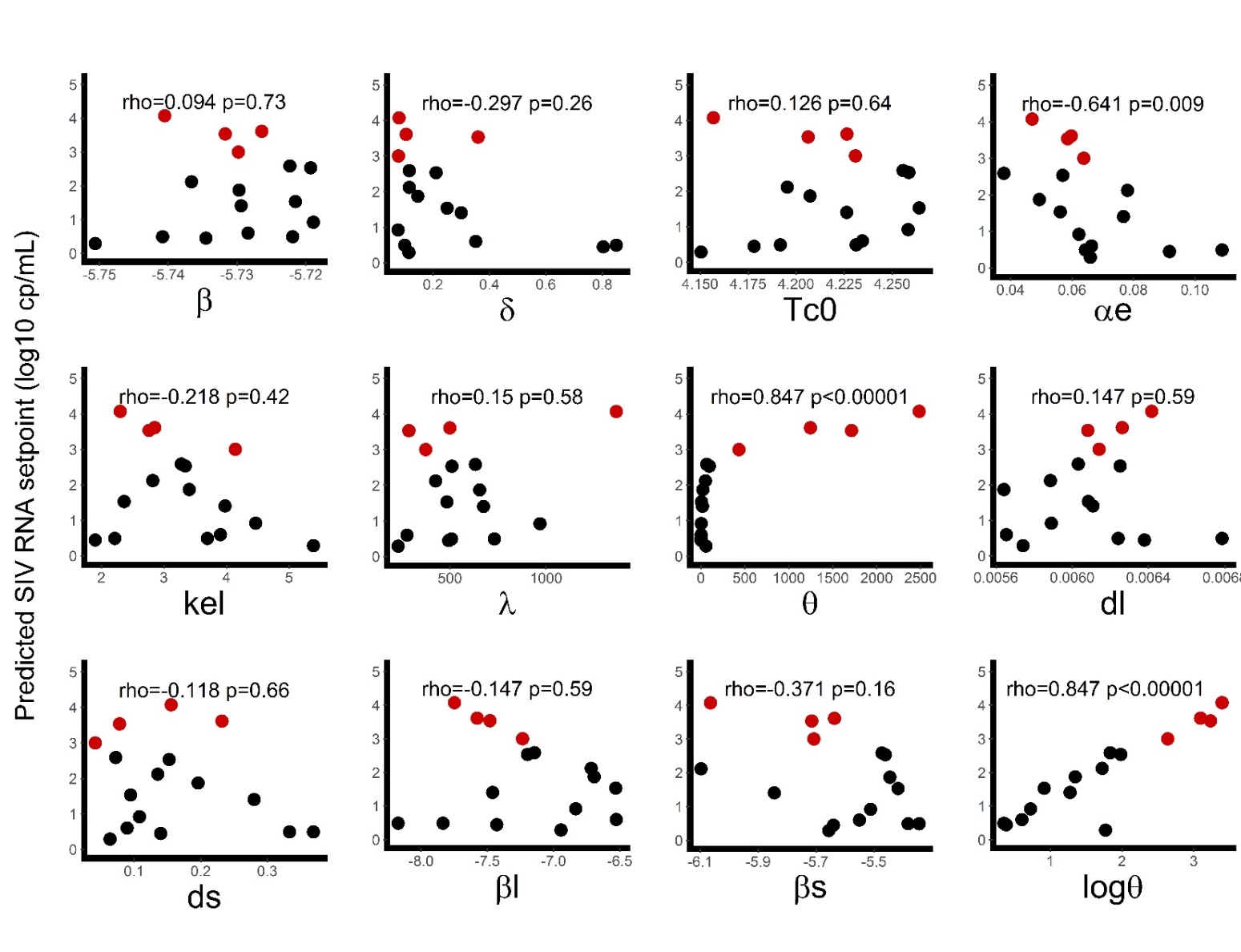

**Figure S5:** Association between individual values of model parameters obtained by empirical Bayesian estimates and individual predicted viral set point. Red dots represent viremic macaques and black dots controller macaques. Corresponding Spearman correlation coefficient and associated p value are reported.

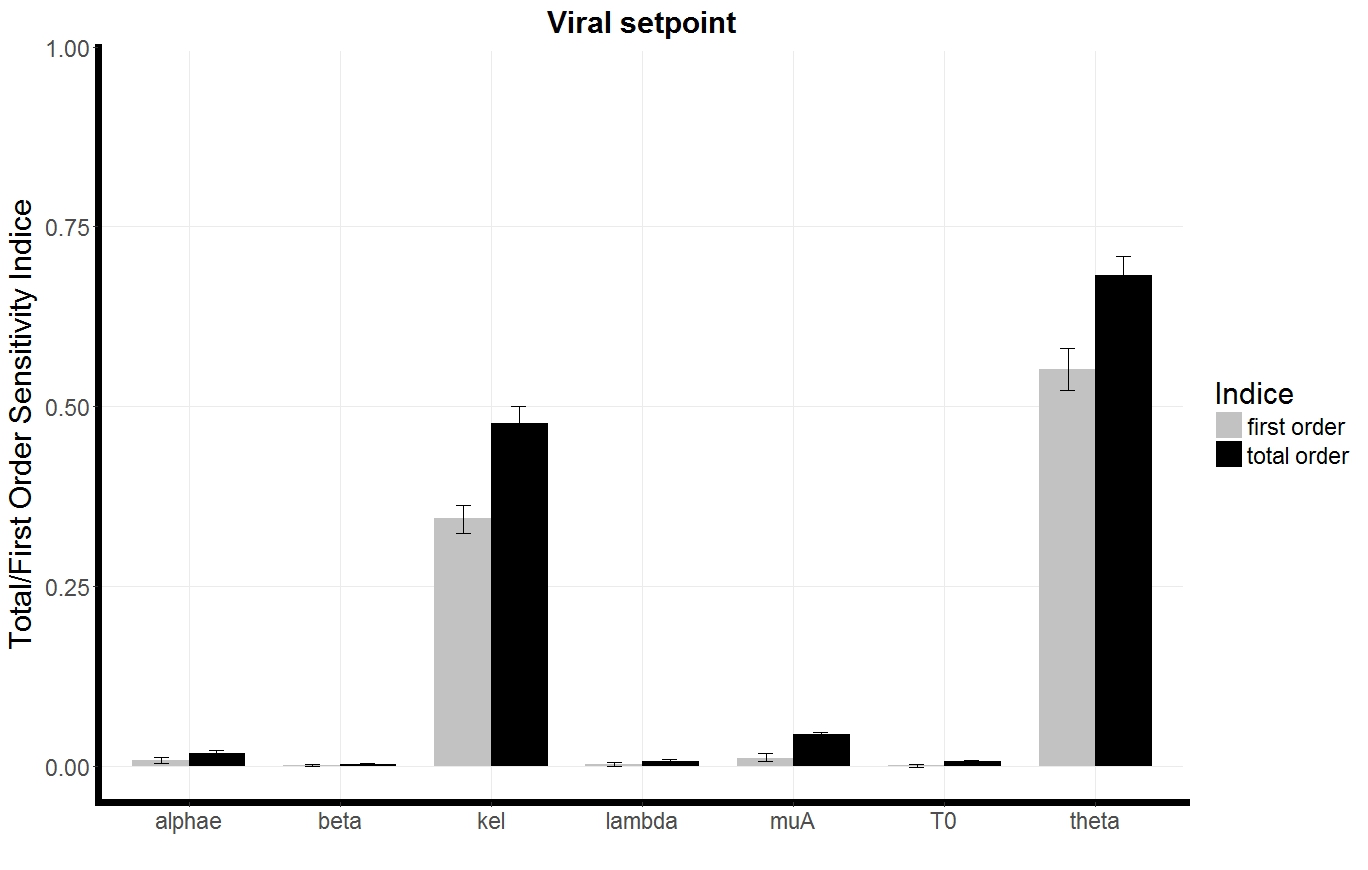

**Figure S6:** First order and total order Sobol sensitivity indices on viral set point for SIV-RNA model parameter. The value of the first order indice represents the impact of parameter variation within the population distribution on the value of the viral set point, with other parameters fixed to their estimated value. The value of the total order quantifies the impact of the parameter variation within its population distribution, when other parameters are allowed to vary within their own distributions.

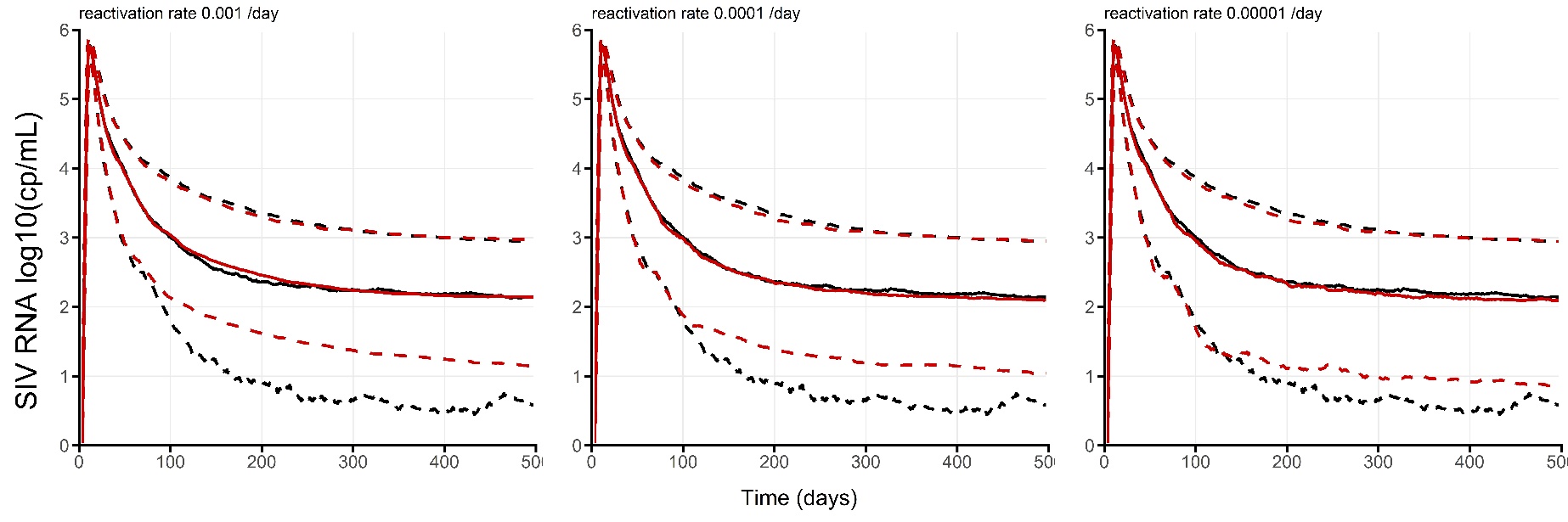

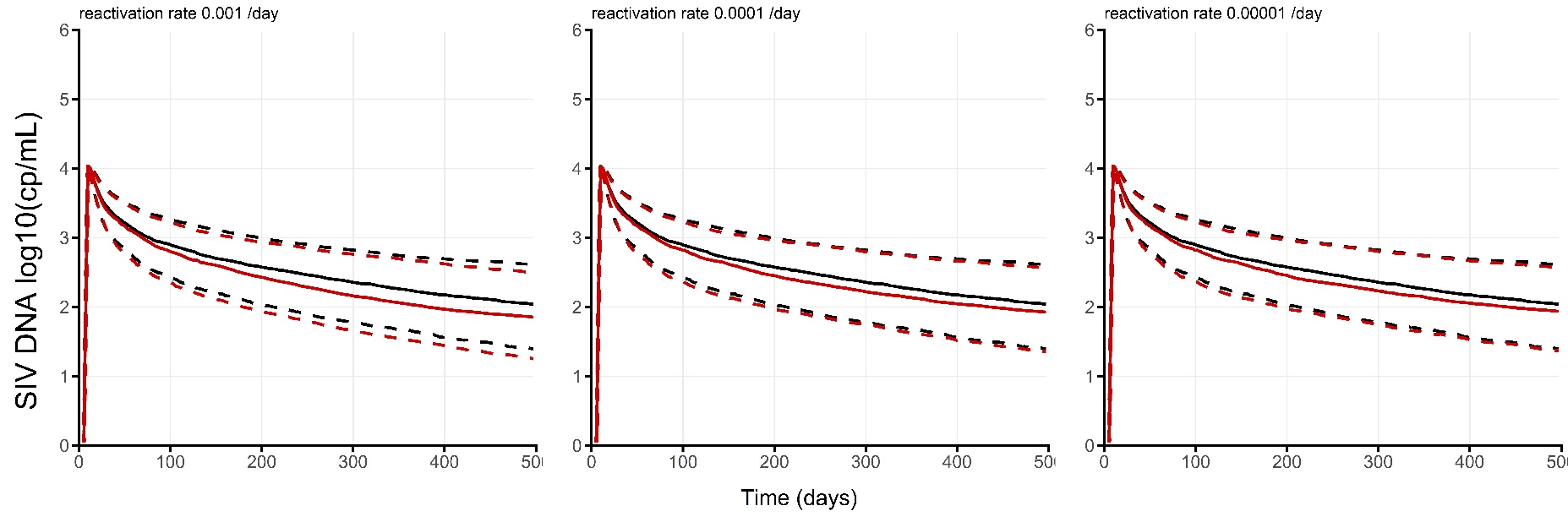

**Figure S7:** Model prediction assuming a reactivation rate of long-lived non-actively producing cells. Median (solid lines) and 25-75 percentiles (dashed lines) of n=1000 simulated SIV-RNA profiles (top panels) and SIV-DNA viral load (bottom panels) for different value of reactivation rate (left: 0.001/day; middle: 0.0001 /day; right: 0.00001 /day). The black lines correspond to the selected model assuming no reactivation.
